# *evosim*: fast and scalable stochastic simulations of evolutionary dynamics

**DOI:** 10.1101/2022.09.28.509950

**Authors:** Dalit Engelhardt, Thomas O. McDonald

**Affiliations:** Department of Data Science, Dana-Farber Cancer Institute, Boston, MA 02215, USA; Center for Cancer Evolution, Dana-Farber Cancer Institute, Boston, MA 02215, USA; Department of Biostatistics, Harvard T. H. Chan School of Public Health, Boston, MA 02115, USA; Department of Stem Cell and Regenerative Biology, Harvard University, Cambridge MA 02138, USA

## Abstract

The simulation of clonal dynamics with branching processes can provide valuable insights into disease progression and treatment optimization, but exact simulation of branching processes via the Stochastic Simulation Algorithm (SSA) is computationally prohibitive at the large population sizes associated with therapeutically-relevant scenarios. *evosim* is a versatile and flexible Python implementation of a fast and unbiased tau-leaping algorithm for the simulation of birth-death-mutation branching processes that is scalable to any population size. Package functionalities support the incorporation and tracking of a sequence of evolutionary changes such as therapeutic interventions as well as the analysis of population diversity. We show that runtimes scale logarithmically with population size, by contrast to the linear scaling of the SSA, and simulations exhibit strong agreement with SSA simulation results. These findings are also supported by mathematical results (Supplementary information).

**Availability:** Package, documentation, and tutorials / usage examples are available on GitHub (https://github.com/daliten/evosim). Mathematical details of the algorithm and the pseudocode are provided in the included Supplementary information.

## 1 Introduction

The dynamical behavior of cell populations is controlled by inherently stochastic processes that determine a wide range of biological outcomes such as gene regulation, evolutionary progression, cellular differentiation, and drug response. Among these processes, births, deaths, and mutations all involve variable degrees of biological randomness that can lead to diverse outcomes under different environmental pressures. The ability to obtain accurate predictions of the interplay between these processes and conditions to which a population of cells is subjected is key to developing better therapeutic interventions, predicting the effects of changing environments on ecosystem dynamics, and understanding the progression of diseases driven by genomic alterations and cell proliferation.

Birth-death-mutation processes, in which a given cell can undergo a division into identical progeny cells, undergo a division resulting in (a) mutant progeny cell(s), or die, are commonly modeled by stochastic branching process models. Gillespie’s Stochastic Simulation Algorithm (SSA) [1, 2] provides an exact simulation of these processes; however, the SSA becomes computationally expensive at large population sizes due to its linear scaling with system size, making the simulation of populations greater than 10^6^ cells typically intractable. This poses significant challenges for biomedical modeling efforts: with a 1 cm^3^ tumor volume containing an ap proximately estimated size of ~10^8^ – 10^9^ cells [3], simulating the evolution and progression of such tumors with the SSA is computationally prohibitive. Even at smaller population sizes (~10^5^–10^6^), studies involving the optimization of treatment or other evolutionary pressures typically require a large number of simulations, which is precluded by the long execution times of the SSA.

As a result, many approximation approaches to the SSA have been developed over the years. Tau-leaping [4], in which multiple events are sampled over the course of a single, longer time interval (leap) underlies a large number of such approaches. Leaping necessarily leads to an efficiency-accuracy tradeoff, and various tau-leaping methods have been developed with the aim of optimizing this tradeoff as well as preventing the population from becoming negative over a single extended leap. These include different approaches to step size selection ([5, 6]), binomial [7, 8]) and multinomial sampling [9], and estimation bias correction [10], among others.

Here, we present a packaged novel tau-leaping algorithm that combines (1) optimized step-size selection, (2) unbiased estimation via event rate rescalings, and (3) distinct ordering in simulation stepping in order to optimize both accuracy and efficiency at any population size and (polyclonal) structure. We show that this approach exhibits high agreement with SSA results while retaining speed and scalability. This approach applies to continuous-time Markov birth-death-mutation branching processes in which a division can result in either two identical progeny cells or one identical and one mutant progency cell (Fig. 1), where cell “types” are defined by their respective rates of birth, death, and mutation to other types. This approach is valid for first-order reaction kinetics in which reactions within species (e.g. birth and death processes) occur at higher rates than interspecies transitions (mutations). Beyond this assumption, our approach is parameter-agnostic in its accuracy and therefore also addresses accuracy gaps in normal-appoximation based approaches [11] that can arise when mutation rates are low.

**Figure 1:**
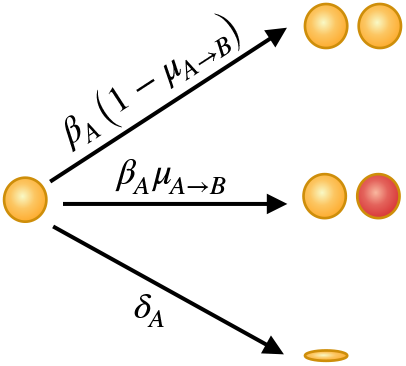
A birth-death-mutation branching process. Cells of type *A* divide at a rate of *β_A_*, die at a rate of *δ_A_*, and experience mutation to type *B* upon division with probability *μ*_*A*→*B*_.

## 2 Description

The *evosim* package is a wrapper for this algorithm and consists of two modes, Simulator and EnsembleSimulator, which can be used for recording a single run (of possibly multiple sequential segments) and an ensemble of simulation runs, respectively. The two simulation modes are designed to permit users to specify a sequence of different conditions experienced by the system over any stretch of time, such as different or alternating treatments that affect the birth and death rates of cells or a serial passaging experiment that dilutes the population when a certain confluence level is reached. An *evosim* simulation run takes as inputs the number of (discrete) cell types (phenotypes, genotypes, or species) that exist or may emerge in the system; the current population distribution, i.e. cell numbers for each of these cell types (which may also be zero for “potential” cell types that could arise by mutation from existing ones); the per-cell birth/division rates and death rates for each cell type; and all probabilities of (inherited) mutations between any two cell types. A cell “type” is uniquely defined by its per-cell birth rate, per-cell death rate, and the probability of mutation from this cell type to all other cell types in the system.

A full simulation may proceed as either a single segment or a sequence of such segments, with different or same values for any of the parameters above, as tracking is maintained until a subsequent reset. This can be used to modify the conditions of the simulation as it proceeds, e.g. introduce a drug through a modification of the birth or death rates. *evosim* comes equipped with tracking and plotting capabilities, a function for subsampling from the entire population (e.g. as in an *in vitro* dilution or surgical resection), as well as functions for computing a number of diversity indices (e.g. Shannon and Gini indices) and for converting the time units on the output; this is particularly useful when the user is interested in long-term evolution (e.g. years) but event rates like births and deaths are specified in short-time units (e.g. days). Optionally, a resource capacity and/or population threshold for terminating the simulation may be specified. Plotting is enabled both for clonal trajectories (i.e. different cell types shown in different colors over time) as well as for total population size over time. Usage examples and tutorials for both the Simulator and EnsembleSimulator modes are provided with the package.

Over the simulation time specified by the user, the simulation proceeds by taking adaptive time leaps that depend on the instantaneous population distribution (subpopulations of all cell types) and the percell birth and death rates of all nonzero-population cell types. The adaptive leaping scheme is based on the observation that small and large populations evolve on different time scales: the expected time to next event (e.g. division) in a small population of cells is longer than the expected time in a large population of cells with the same per-cell division rate. By assigning an “effective” time scale to the simulation (see mathematical details in the Supplemental Information), the simulation proceeds with logarithmic time complexity 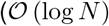 in total population size *N*) rather than linear complexity 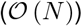, thus permitting scalability (Fig. 2). Since, as with any tau-leaping scheme, these leaps are larger than the fundamental time scale of the system, a naive application would lead to a biased estimation of the system dynamics. By rescaling the birth and death rates through a derived transformation (see the SI), this bias is removed from the simulation, resulting in very high agreement between the tau leaping simulation results and those of the SSA. Additionally, within each leap populations are sampled in the order of their effective time scales (see pseudocode and explanation in the SI) in order to ensure highly accurate modeling of the effect of mutations on population growth at any population scale without simulation efficiency loss. We note that due to a bias-variance tradeoff, unbiased tau-leaping results in a slight loss of variance (see SI for mathematical results); however, this is a second-order effect that in practice carries minor effect (Fig. 2).

**Figure 2:**
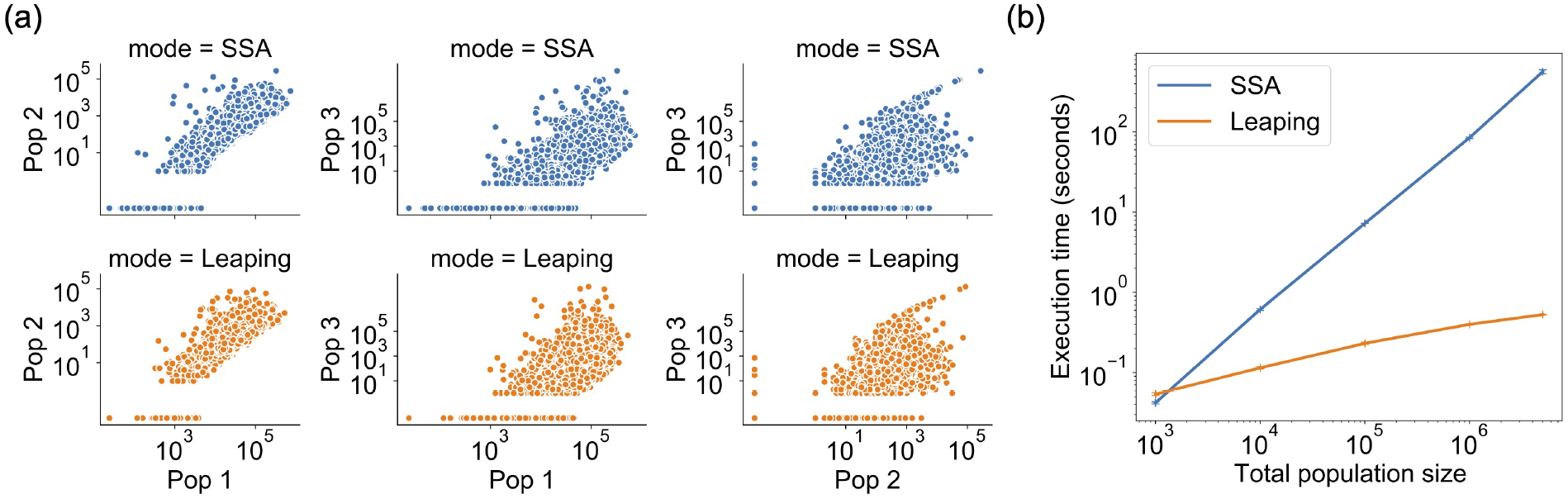
Comparison of *evosim* performance using its tau-leaping algorithm versus the SSA for a three-type simulation starting from a single cell. (a) Projections of 5000 simulations for each pairwise combination of cell types shows strong agreement in the clonal distributions between the leaping algorithm and the SSA; (b) Execution times for 30 simulations (same setup as (a)) up to different total population sizes show a logarithmic increase with total population size when tauleaping is used and an exponential increase with total population size for the SSA. Parameters used: *β*_1_ = *β*_2_ = 0.004 cells/day, *β*_3_ = 2*β*_1_, *δ_i_* = 0.1*β*_1_ for *i* = 1, 2, 3, *μ*_1→2_ = 10^−3^, *μ*_1→3_ = *μ*_2→3_ = 10^−4^. Initial population sizes were set at *n*_1_ = 1, *n*_2_ = *n*_3_ = 0. Simulations were set to terminate after 8.5 years of simulation time. For plotting on a logarithmic scale, *ϵ* = 0.01 was added to results with zero cells.

## 3 Results

A branching process of single cell type with no death takes an average of 212 milliseconds up to a population size of 10^9^ and 300 ms up to 10^12^ cells on a personal 2.7 GHz Intel Core i7 CPU with 16 GB RAM running a Jupyter notebook server in OS X. Including a mutation probability per division of 0.01 (two-type process, each type starting at 1 cell) results in an average runtime of 602 ms up to 10^9^ cells and 879 ms up to 10^12^ cells. Simulation results confirming high agreement between leaping simulations and the SSA are shown in Fig. 2 and in the SI.

## Acknowledgements

We are grateful to Franziska Michor and the Michor Lab for helpful discussions.

## Funding

This work was supported by the Helen Gurley Brown Presidential Initiative at Dana-Farber Cancer Institute and the Center for Cancer Evolution at Dana-Farber Cancer Institute. Research reported in this publication was supported in part by The National Institute of General Medical Sciences of the National Institutes of Health under grant number R01GM144962.

## Supplemental Information

### 1 Introduction

We present here the mathematical and computational details of the tau-leaping algorithm implemented in the evosim package. In Section 2 we set out notation and define the concept of *characteristic-time leaping*, a simulation algorithm that incorporates tau-leaping but sets the leap size to the expected time to next event. In Section 3 we present our method (*extended-time leaping*) for extending these leaps and accordingly rescaling the sampling probabilities to retain simulation accuracy. In Section 4 we derive the efficiency gain of extended-time leaping over the SSA for the single-species case. Supporting simulation results and pseudocode are included.

### 2 Preliminaries

#### 2.1 Notation and basic tau-leaping

We consider *C* interacting cell types, i.e. such that transitions between different cell types may occur. The state of the system at time *t* is described by **x**(*t*) = (*x*_1_(*t*), …, *x_C_*(*t*)) where *x_j_*(*t*) is the population at time *t* of the *j*th species. The set of self-interactions and inter-species transitions of species *j* are described by size-dependent rates {*r_j_* (*x_j_*)}, *j* ∈ {1, …, *C*}.

Gillespie’s Stochastic Simulation Algorithm (SSA) [1, 2] is based on the assumption that the time *τ* to the next event in this system is an exponentially-distributed random variable with mean 1/∑_*j*_ *r_j_* (*x_j_*). The interval in which *χ*_2_ is found, SSA thus proceeds bLy generating two random numbers *χ*_1_, *χ*_2_ ∈ [0, 1]; the time to next reaction is computed as τ = log (1/η_1_)/∑_*i*_ *r_i_* (*x_i_*), and the next event is chosen in accordance with the reaction probability interval in which *χ*_2_ is found, where the event probability for event type *j* is given by *r_j_* (*x_j_*) /∑_*i*_ *r_i_* (*x_i_*). The populations are then updated according to the chosen reaction. Since this procedure quickly becomes very computationally expensive at larger system sizes, a number of methods have been developed to increase simulation efficiency. The focus of this work will be on tau-leaping, whose basic form assumes that an interval *τ* can be selected such that the change in **x**(*t*) over *τ* does not lead to appreciable changes in any of the rate functions. We denote by **K**_*j*_ (*τ*; **x**, *t*) the number of events, i.e. transitions or self-interactions, 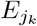 (where *k* indexes different events pertaining to species *j*, e.g. birth, death, or mutation to a different type), taking place within [*t, t* + *τ*] in species *j*. We define 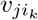 as the change in *x_j_* due to an event 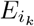, where *i_k_* indexes the set of events ℰ_*i*_ involving species *i*. The state of the system at *t* + *τ* is updated as

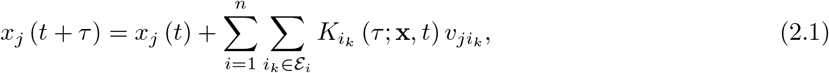

where 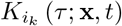 are random numbers drawn from Poisson distributions with means 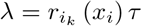.

We will restrict ourselves to rate functions *r_j_* whose size-dependence is given by *r_j_* (*x_j_*) = *g_j_x_j_*, where *g_j_* is a fixed constant that defines the event rate per individual in cell type *j* (e.g. the rate at which a single cell divides or dies)^1^. Since we consider here branching processes, the system will consist of cell types *X_j_*, *j* ∈ (1, ․, *C*) undergoing births, deaths, and mutations via the following reactions

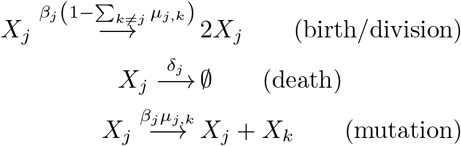

occurring with respective rates *β_j_* (the rate at which a single cell of type *j* divides), *δ_j_* (the rate at which a single cell of type *j* dies), and *β_j_μ_j,k_* (the rate at which a single cell of type *j* divides into a *j* cell and a mutant *k* cell), where *μ_j,k_* is the probability of mutation from type *j* to type *k*. As noted in the main text, we will assume throughout that *μ_j,k_* ≪ 1 for all *j*, *k* (this is equivalent to the assumption that mutations occur at much lower rates than birth and death rates). When referring generically to any of these rates, we will use the notation *g_j_*, where

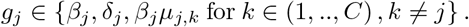

For simplicity of notation in subsequent sections, we will use in parts of the derivation the notation *g_j,±_* for *β_j_*(*g_j,_*_+_) and *δ_j_* (*g_j,−_*).

#### 2.2 Characteristic-time leaping

The probability of an event *E* with rate *r_j_* (*x_j_*) = *g_j_x_j_* occurring at some point over the interval *τ* is given by

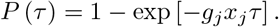

When *τ* = 1/(*g_j_x_j_*), in expectation a single event *E* will occur over the interval *τ*. We accordingly define the *characteristic time τ*_char,*j*_ of cell type *j* at time *t* as

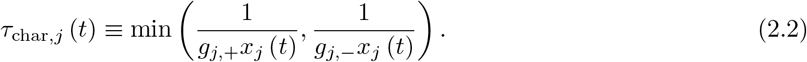

This is the expected time to the first event (single member addition or removal) in the population of cell type *j*. Simulating the dynamics of cell type *j* by tau-leaping on intervals of length *τ*_char,*j*_ (*t*) is a close approximation to exact simulation, as only one event is expected in each interval. i.e.

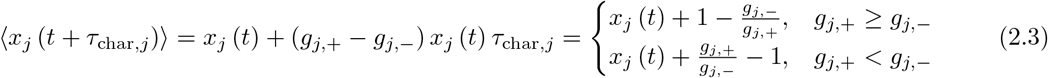

where we have incorporated the expectation values of Poisson draws *λ_j_* = *g_j_x_j_τ* (see Eq.2.1). For simplicity, recalling that by assumption *μ_j,k_* ≪ 1 ∀ *j*, *i*, we have neglected the contribution of *μ_j,i_* to the rates, which would lead to an adjustment *g*_*j,*+_ → *g*_*j*,+_ (1 − ∑_*i*≠*j*_ *μ_j,i_*). If a simulation is carried out over *m* such steps, then in expectation (where at each step we consider the expectation)

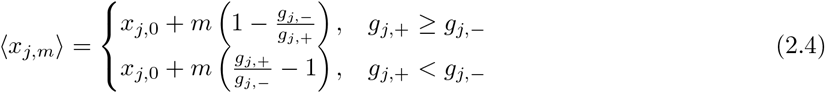

where *x_j,m_* is the population of cell type *j* at the beginning of the *m*-th step, so that *x*_*j*,0_ = *x_j_* (*t*) for a simulation interval staring at time *t*. We will refer to each step *m* → *m* + 1 of such a simulation as *characteristic-time leaping*. We emphasize that the expectation value of the change in the population of *j* is constant in each such step, as indicated in (2.4), but that the time interval corresponding to each such step (*τ*_char,*j*_) will vary with the population size (2.2) and hence may change with each step. Written differently, over an interval of length *T* we have in expectation that

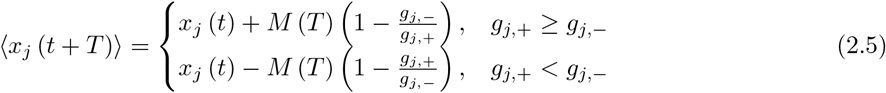

where *M* (*T*) is the number of *τ*_char,*j*_-length simulations (each of potentially different length), i.e. the number of Poisson sampling draws for each event, that cover the interval *T*. In expectation, *M* is the solution to

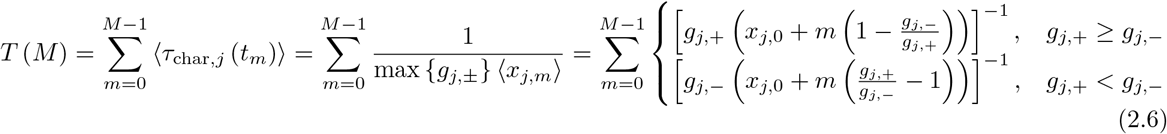

where (see Eq. 2.7)

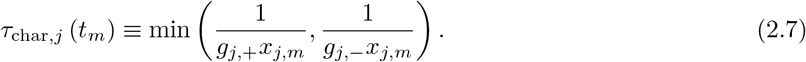

In closed form (2.6) is given by

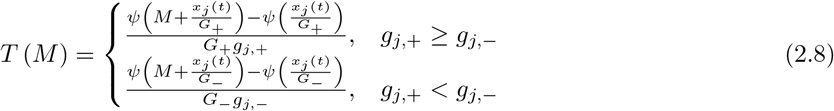

where *ψ*(*z*) = Γ′ (*z*) /Γ (*z*) is the digamma function and, for notational simplicity, we have defined

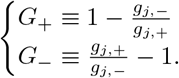

#### 2.3 Multinomial sampling

As with the SSA, characteristic-time leaping scales with population size and becomes computationally expensive at large populations due to the large number of sampling draws required. We wish to establish whether leaping on longer intervals can sufficiently well approximate characteristic-time leaping; however, the possibility of negative population counts is a general feature of tau-leaping while sampling from a Poisson distribution since at larger values of *M* the population may become negative in a single leap. One approach for preventing negative population counts during tau-leaping is to substitute Poisson sampling for binomial sampling [3, 4]. In binomial tau-leaping the number of events is sampled from a binomial distribution with mean *n_j_p_j_* = *λ_j_* = *g_j_x_j_τ*, where *λ_j_* is the Poisson parameter, under the assumption that this provides a good approximation in the regime of large *n_j_* and small *p_j_*, where *n_j_* is the current population size of species *j* and *p_j_* is the probability of having the event of interest (e.g. a cell division) occur in any member of species *j*. Here, since a cell type can have multiple rates *g_j_*, we will consider the multinomial distribution. We will first establish the validity of multinomial sampling in the case of characteristic-time leaping – where the characteristic-time Poisson sampling described above is the baseline for comparison – before proceeding to the case of extended leap times. We will first focus on the rates *g*_*j*,±_ that add or remove members of cell type *j* and return to inter-type transition rates (i.e. mutations) in Sec 3.2.

When sampling from the multinomial distribution we consider the set of probabilities *p*_±_ (*τ*) = *g*_*j*,±_*τ* as well as the probability that no change has occurred, *p*_nc_ (*τ*) = 1 − *p*_+_ (*τ*) − *p*_−_ (*τ*), so that the expected size of the population after a sampling is given by

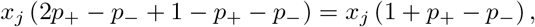

where the first two terms represent, respectively, cell division (hence the factor of 2) and cell death. Then, substituting *p*_±_ = *g*_*j*,±_*τ*, we observe that

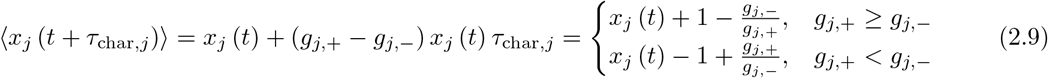

identically to that obtained through Poisson sampling above (2.3), showing that multinomial sampling results in an unbiased estimate of the population size (as compared to Poisson) in characteristic time simulations. We can show (A.3) that the variance is given by

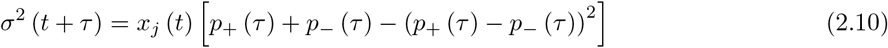

or, once again substituting *p*_±_ = *g*_*j*,±_*τ*,

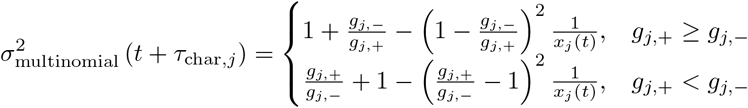

The variance in Poisson sampling will be the sum of variances *λ*_*j*+_ and *λ*_*j*−_,

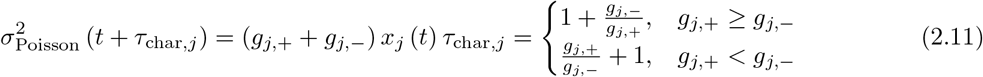

so that the fractional deviation in the variance is of 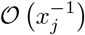:

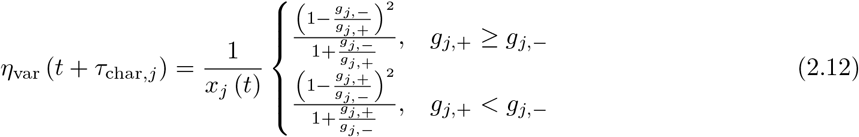

and hence at large *x_j_*, as expected, multinomial sampling provides an excellent approximation to Poisson sampling. We will define *x*_thresh_ at the threshold value at which we consider (2.12) to be acceptably low (e.g. *x_j_* ∼ 100 − 1000).

### 3 Extended-time leaping: estimation accuracy

#### 3.1 Self-interactions

Let 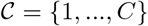 be the set of all cell types and define

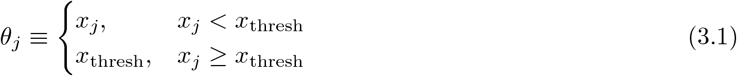

and the *effective* characteristic time

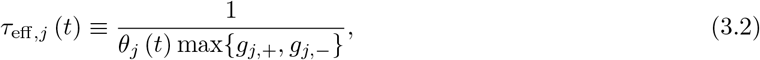

while recalling that the true characteristic time is given by

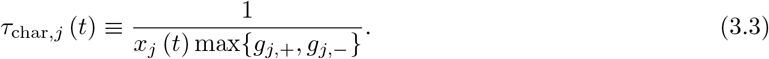

Then we set the simulation time step of the system at time *t* to

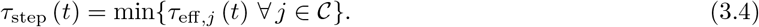

We will refer to simulations that involve a single draw from a multinomial distribution Mult (*n_j_*, **p**_*j*_), where **p**_*j*_ = (*p*_*j*,+_, *p*_*j*,−_, 1 − *p*_*j*,+_ − *p*_*j*,−_) over the time interval *T* = *τ*_step_ (*t*) as *extended-time leaping*.

We define the following transformations of the multinomial sampling parameters:

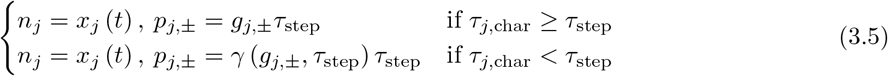

where

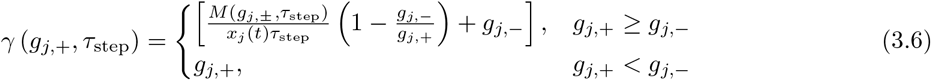

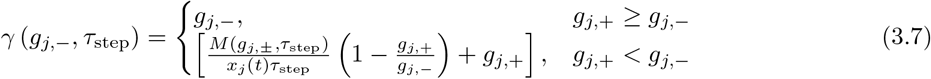

such that *M* (*g*_*j*,±_, *τ*_step_) is the solution to (2.8) with *T* = *τ*_step_.

##### Claim 1.

*Under the transformations of* *Eqns. 3.5*, *3.6*, *and* *3.7*, *extended-time leaping of leap size T* = *τ*_step_ *does not lead to any appreciable degradation in the estimation accuracy of the mean and variance of the resulting population distributions for any cell type* 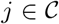 *as compared with characteristic-time leaping (with characteristic time τ*_char,*j*_ *over the interval τ*_step_.

*Proof.* If for cell type *j τ*_char,*j*_ > *τ*_step_, multinomial sampling at more frequent intervals simply represents a closer approximation to the continuous-time simulation limit, and therefore no loss of accuracy is incurred. If *τ*_char,*j*_ < *τ*_step_, we have that under multinomial sampling the expected value will be given by

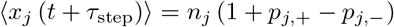

so that, substituting in the probabilities *p*_*j*,±_ (3.5, 3.6, 3.7), we have

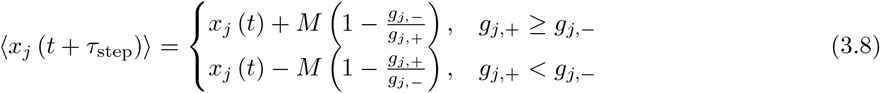

where *M* = *M* (*g*_*j*,±_, *τ*_step_). From (2.5), , where *T* = *τ*_step_, we see that the expected value (3.8) after an extended *τ*_step_ leap is the same as that obtained over the requisite number of characteristic-time Poisson sampling steps.

The variance is given by (2.10):

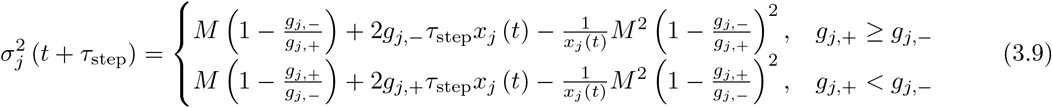

To see how (3.9) compares to the variance from characteristic-time leaping, we note that when *x_j_* (*t*) is large,i.e. *x_j_* ≥ *x*_thresh_, as we assume here, we can asymptotically expand the digamma functions *ψ* (*z*) ∼ log *z* in (2.8) to obtain the solutions

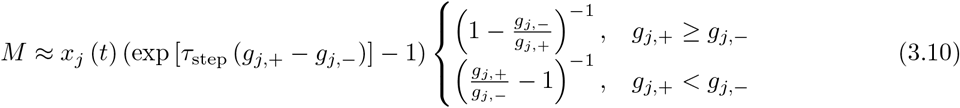

Next, we let *α* denote the species for which *τ*_eff*,α*_ = *τ*_step_ (*α* may or may not be the same as *j*) and further expand (3.10) in *x*_thresh_ (recalling the expression for *τ*_step_, Eqns. 3.2 and 3.4)) as

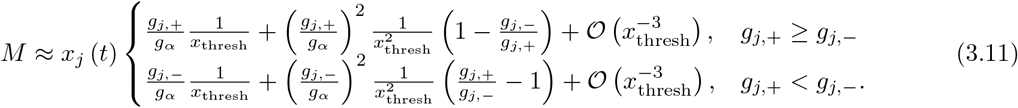

Substituting into (3.9), we obtain

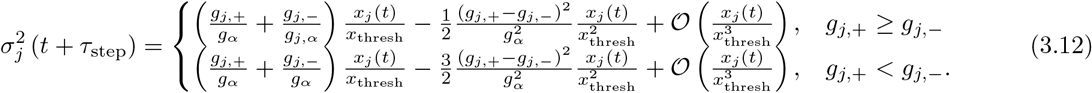

On the other hand, *M* (*τ*_step_) steps of characteristic-time Poisson simulations result in a cumulative variance (see Eq. 2.11) of

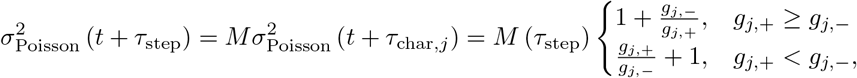

which, substituting *M* from the expressions (3.11), becomes

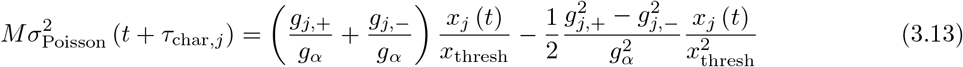

We see that to leading order the variance obtained in *M* (*τ*_step_) Poisson characteristic-time simulations (3.13) is the same as that of a single leap *τ*_step_ with multinomial sampling (3.12), and the fractional deviation in variances is of 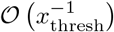. This deviation is on the same order as that obtained in (2.12) for characteristic-time leaping, which we considered to be acceptably low for a fixed choice of *x*_thresh_ ∼ 100, with extended-time leaping slightly overrestimating the variance (by, to leading order, 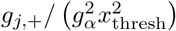) in the population of *j* if *g*_*j*,+_ > *g*_*j*,−_ or if *g*_*j*,−_ > 2*g*_*j*,+_ and slightly underestimating the variance if *g*_*j*,+_ < *g*_*j*,−_ < 2*g*_*j*,+_.

Fig. 1 shows a comparison of single cell type simulation outcomes for extended-time leaping and the SSA for two different leaping threshold levels (*x*_thresh_ = 10^2^, 10^3^) over three different simulation time intervals, with identical initialization parameters used for all simulations. Strong overlap in population sizes is observed in all cases. SSA runtimes are shown to increase linearly with population size whereas extended-time leaping runtimes only increase logarithmically with population size.

Finally, we note that the transformations (3.7) and (3.6) amount to transformations of the demographic rates. I.e. we have shown that extending the simulation interval while implementing an appropriate rescaling of the relevant event rates results in a similar distribution of outcomes (unbiased estimate, with an acceptably low deviation in variances). Substituting in the expression for *M* (3.10), we can approximate the rate transformations as

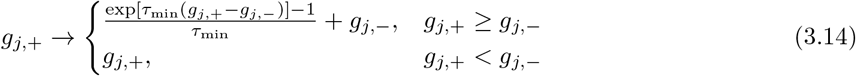

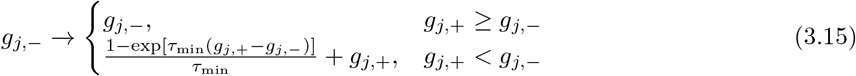

If we expand to second order, we see that the rate transformations are given by

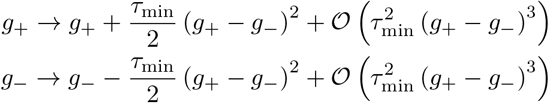

so that extended-time leaping retains simulation accuracy while simulating in larger steps by increasing birth rates (*g*_*j*,+_) when a subpopulation is growing (*g*_*j*,+_ > *g*_*j*,−_) or decreasing death rates (*g*_*j*,−_) when a subpopulation is shrinking (*g*_*j*,+_ < *g*_*j*,−_).

#### 3.2 Inter-species transitions

While rate rescaling in *g*_±_, as described in the previous section, ensures an unbiased estimate of the growing or declining species subpopulation, we must further ensure that flux to other subpopulations is also estimated correctly under extended-time leaping. For a transition (mutation) probability *μ* ≡ *μ_j,k_* from subpopulation *j* to subpopulation *k*, the expected number Ω_*j*→*k*,ETL_ of new members of *k* as a result of transitions from *j* after time interval *τ*_min_, starting at time *t*_0_, is given by *μx_j_* (*t*) *p*_*j*,+_ (*τ*_step_). From (3.5) and (3.6), this is given by

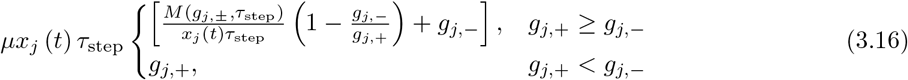

**Figure 1:**
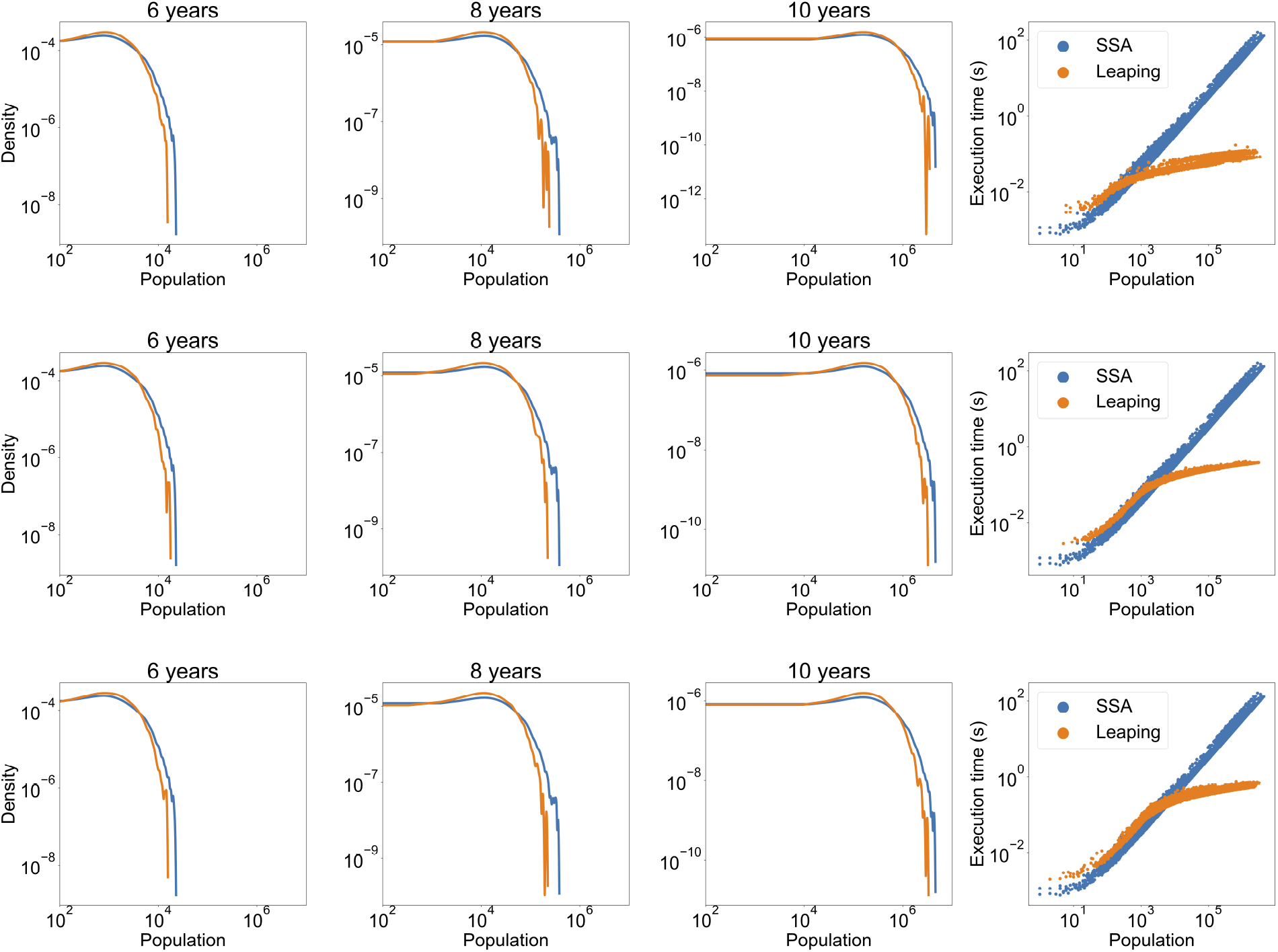
Single-species simulations show strong agreement and a high efficiency increase factor between extended-time leaping (orange) and the SSA (blue). In each panel the first three plots show the population size density distribution over 5000 SSA and 5000 extended-time leaping simulations for the specified number of simulated years and *x*_thresh_, and the rightmost plot shows the respective (leaping and SSA) distributions of execution times versus population size. Simulation parameters were set at *g*_+_ = 0.004 day^−1^, *g*_−_ = 0.1*g*_+_ and initiated with a single cell in all cases. All plots are shown on a log-log scale.

On the other hand, after *M* steps of characteristic-time leaping, in expectation the total flux to *k*, Ω_*j*→*k*,CTL_, is given by

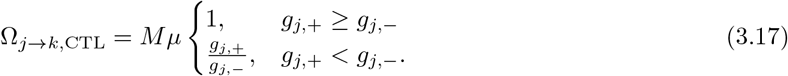

We can therefore show that a rescaling of the mutation rate *μ* → *μ^′^* such that

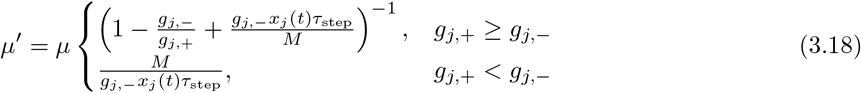

will result in equality of expected fluxes under extended-time leaping.

As we loop through sampling the different subpopulations (see Algorithm 1), mutations are added to the subpopulations into which they occur. Since intervals are larger in the leaping regime than the characteristic time scales of system, there is a potential risk of underestimating the effects of mutations on smaller populations: once a cell has been born in a certain cell type, that cell can then divide and thus continue to expand the this subpopulation; however, if such mutations are only accounted for after many characteristic time steps, the subpopulation size will be underestimated. To counteract this effect, at each simulation step all cell types are ordered by their effective characteristic times, and sampling is performed in this order. As a result, populations with larger time scales that evolve more slowly receive their proper mutation influx within their characteristic time step, hence avoiding the underestimation problem (see Algorithm 1).

##### Algorithm 1: Extended-time leaping for subpopulations *j* ∈ sim_pops over time interval sim_time

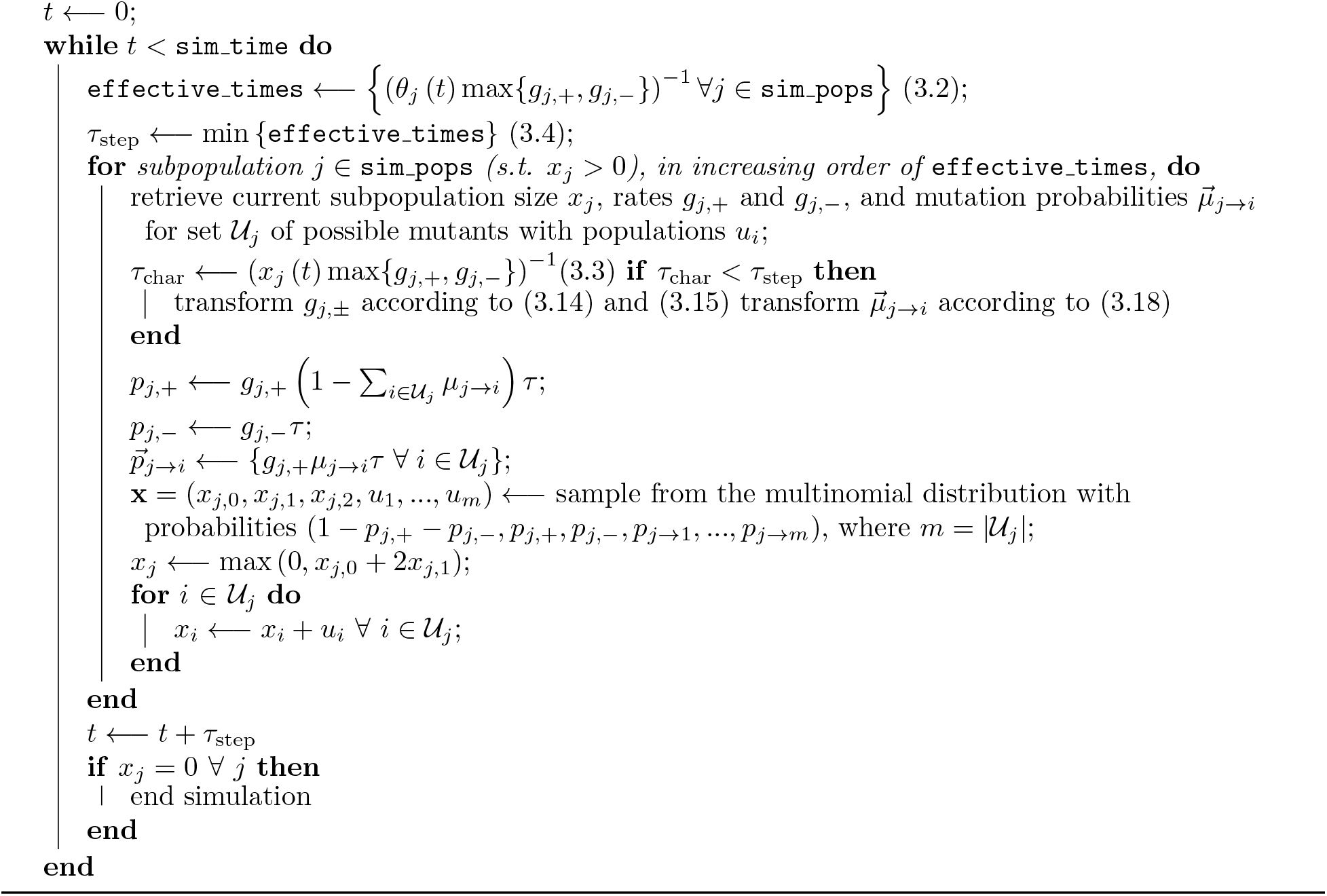

Fig. 2 shows a comparison of how well extended-time leaping performs against the SSA in two-species simulation when rescalings of the sampling probabilities are implemented versus when extended-time leaping is carried naively without any rescalings. Both simulation setups shown in Fig. 2 exhibit high agreement with the SSA when rescaling is implemented and decreased agreement otherwise.

### 4 Extended-time leaping: time complexity

#### 4.1 The single-species case

The time complexity of a population growth simulation is defined by the number of steps – each consisting of a multinomial sampling – needed to grow its population from an initial size *n*_0_ to a final specified size *N*. We first consider the case of a single cell type with *g*_+_ > *g_−_*. When *n*_0_, *N* < *x*_thresh_, then the simulation is in the characteristic-leaping regime for its entire duration. In this case, from (2.5),

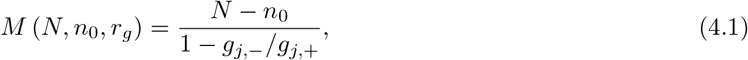

**Figure 2:**
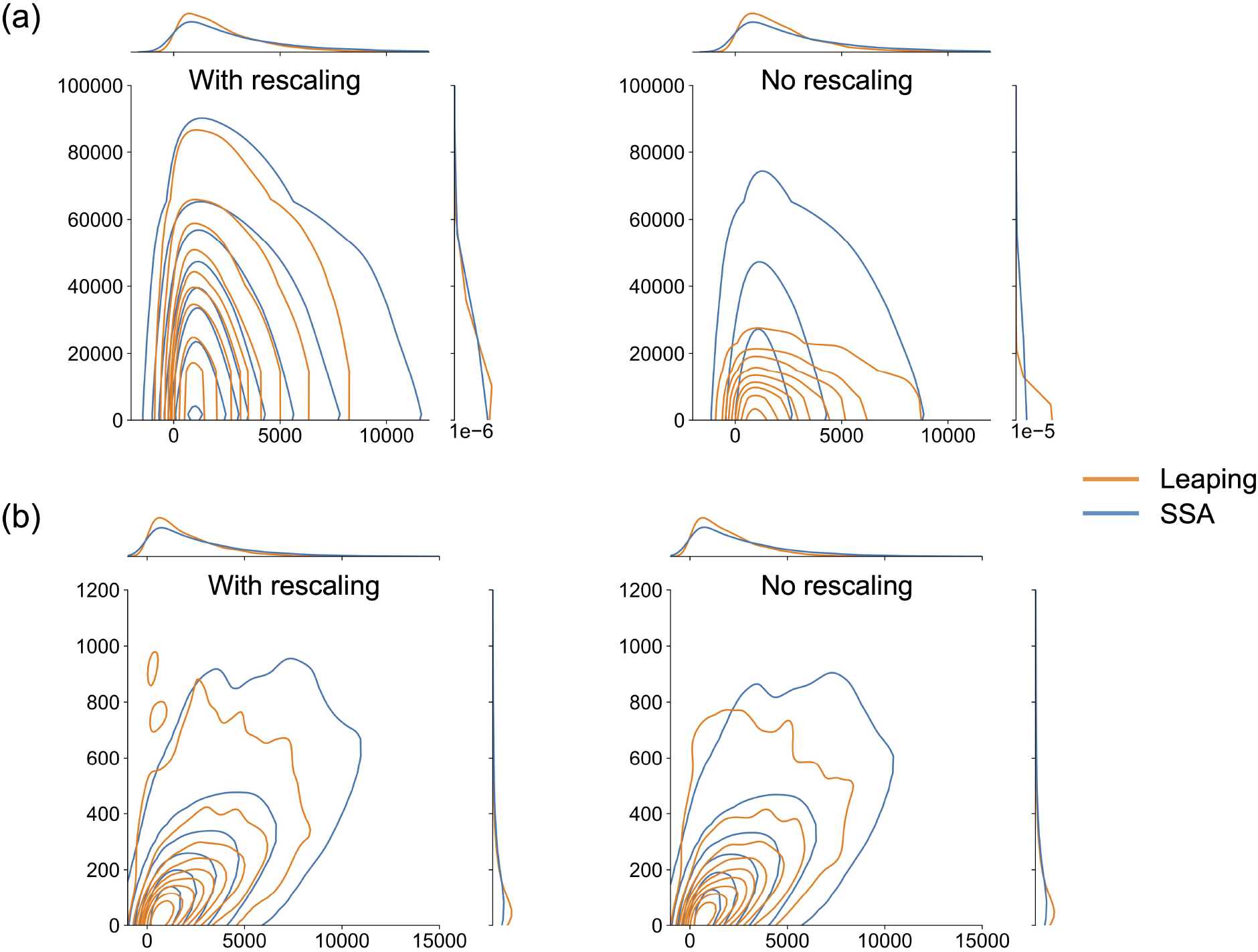
Joint distribution KDE plots of the population distribution for two-species simulations show strong outcome agreement between extended-time leaping (orange) for the default *x*_thresh_ = 10^2^ and the SSA (blue) when rescaling is implemented (left column plots) but lower agreement when leaping is done without rescaling (right column plot), where (a) and (b) represent different evolution initialization parameters. The extent of significance of rate rescaling is higher in (a) than in (b) but evident in both cases. Initialization parameters: (a) *β*_1_ = 0.004/day, *β*_2_ = 0.008/day, *δ_i_* = 0.1*β_i_* for *i* = 1, 2, *μ*_1,2_ = *μ*_2,1_ = 10^−3^; *δ_i_* = 0.1*β_i_* for *i* = 1, 2, *μ*_1,2_ = *μ*2, 1 = 10^−2^; (b) *β*_1_ = *β*_2_ = 0.004 day^−1^, *δ_i_* = 0.1*β_i_* for *i* = 1, 2, *μ*_1,2_ = *μ*_2,1_ = 10^−2^. Both cases were initiated with a distribution of *x*_1_ = 1, *x*_2_ = 0 and run for a simulation time of 6 years; distributions represent outcome from 5000 simulations in each case.

steps are taken to reach size *N*. i.e. characteristic-time leaping is, as expected, 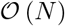 complexity. If no extended-time leaping is implemented, the simulation is of linear time complexity for arbitrarily large *N*.

With extended leaping, *τ*_step_ = 1/ (*x*_thresh_*g*_+_), and every leap *ℓ* covers a number *M_ℓ_*; of characteristic-time steps (see Eqn. 2.6):,

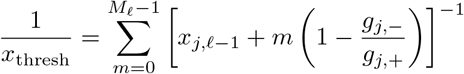

where *x*_*j*,ℓ_ is the population size at the end of leap *ℓ* − 1, i.e., recalling (2.8),

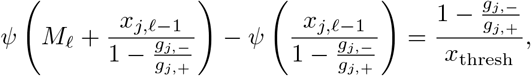

or, again using the digamma asymptotic expansion,

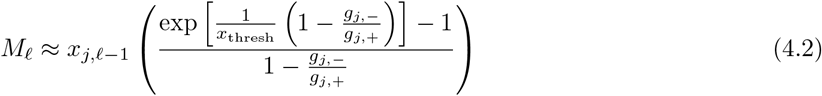

where, as we have previously established,

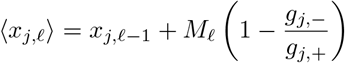

which we can therefore write as

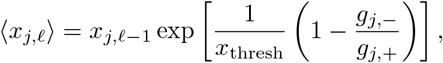

and which, in expectation, has the explicit solution

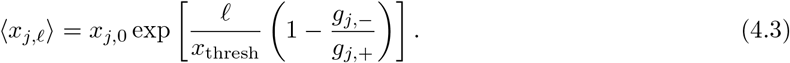

Recall that *M* constitutes the number of multinomial samplings under characteristic-time leaping. We denote by *L* the number of multinomial samplings under extended-time leaping, i.e. *L* represents the number of extended-time leaps (of length *τ*_min_). When leaping starts at population *n*_0_, then, from (4.3),

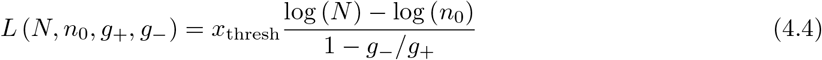

samplings will be needed by the time the population reaches size *N*, i.e. extended-time leaping is 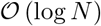 complexity.

In a one-species SSA simulation (see Algorithm 2), on the other hand, at the *m*-th simulation step the following may occur with probabilities *p*_±,0_:

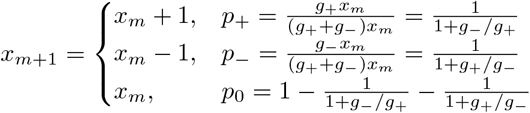

so that

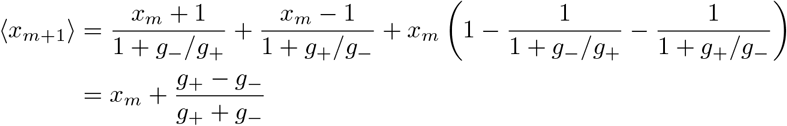

In expectation, then, after *M* steps, starting from population *n*_0_, we have that

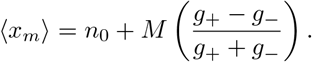

For a final population *N*, we can solve for *M* as

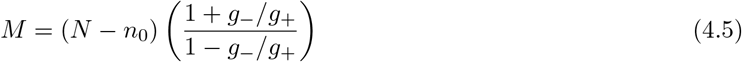

so that the SSA operates with linear, 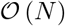 complexity.

The efficiency gain to extended-time leaping with threshold *x*_thresh_ from *x* = *x*_thresh_ onwards is given by

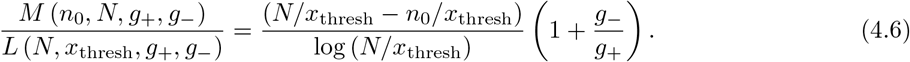

Fig. 3 shows a comparison of speed and number of simulation steps for a SSA simulation versus extended-time leaping set at multiple thresholds. Simulation results confirm the logarithmic complexity of extended-time leaping versus the linear complexity of the SSA. We show in Fig. 4 as well as in the main text that these relationships continue to hold for larger numbers of cell types.

**Figure 3:**
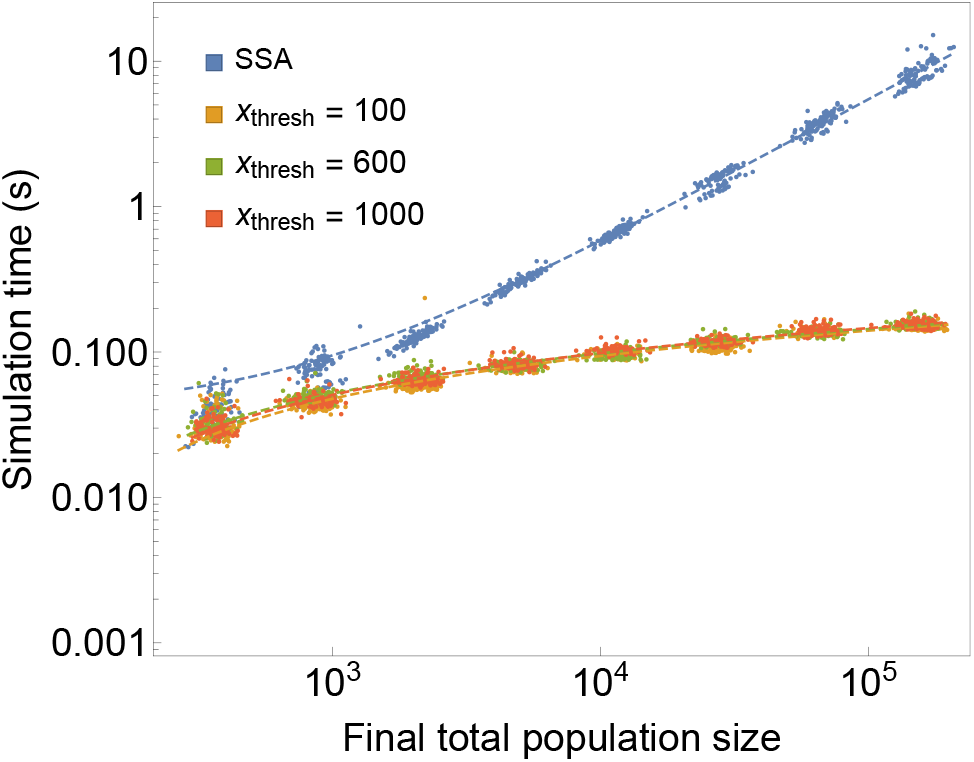
Extended-time leaping leads to significant gains in simulation speed. Shown above is total simulation time versus the population size at simulation termination for single-species growth simulations. All simulations were started at population size *x* = 100 and terminated at a sequence of increasing times (30 through 170 months, in 20 month intervals, with 100 simulation runs at each termination point). The fits shown in dashed lines represent logarithmic fits (Eqn. 4.4) for population average versus execution time (note that Eqn․ 4.4 is representative of number of steps, of which runtime is a proxy) at each termination point for *x*_thresh_ = 100 (*r*^2^ = 0.982), *x*_thresh_ = 600 (*r*^2^ = 0.987), and *x*_thresh_ = 1000 (*r*^2^ = 0.995), and a linear fit (Eqn. 4.5) for the runtime and population averages for the SSA (*r*^2^ = 0.9997). Simulation parameters used were the same as in Fig. 1.

**Figure 4:**
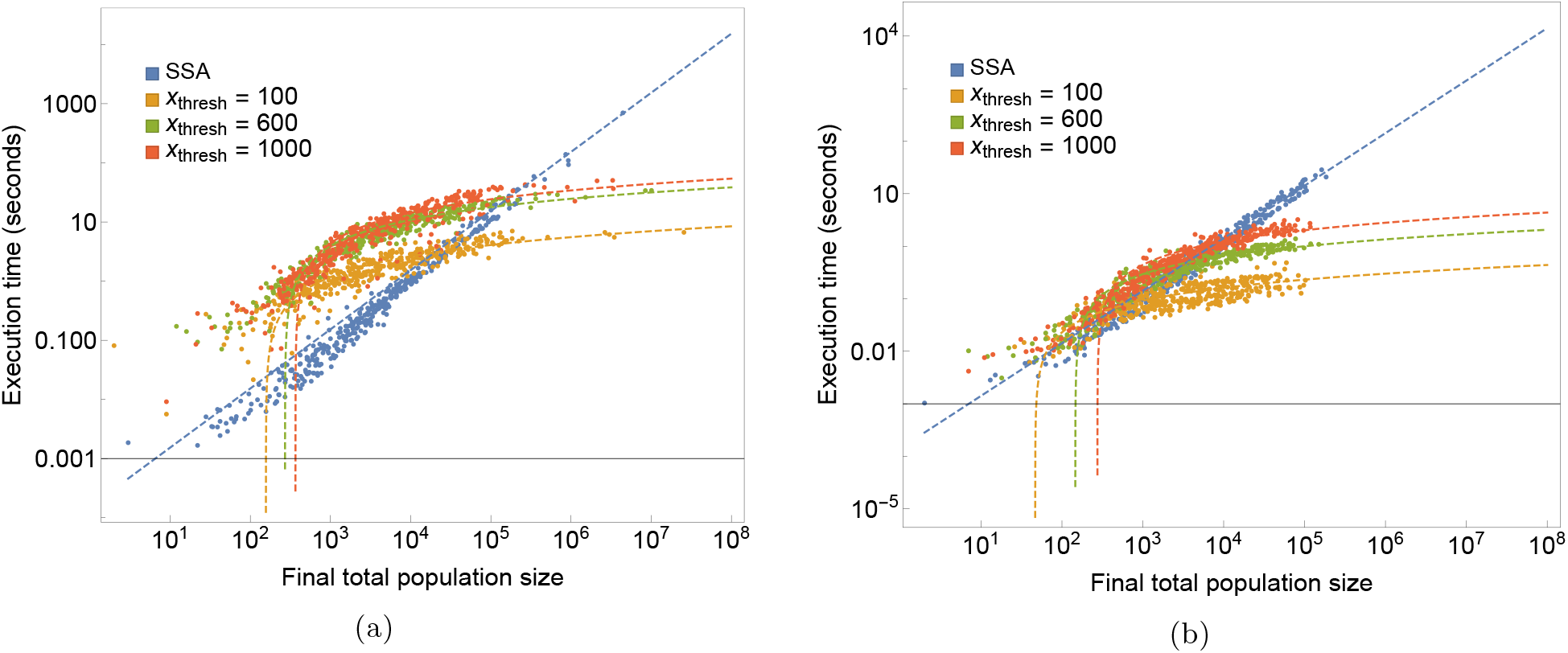
Execution times scale linearly with population size for the SSA and logarithmically with population size for extended-time leaping. Figures (a) and (b) show, respectively, the execution times versus total population size for the configurations shown in (a) and (b) of Fig. 2 at different leaping thresholds. Simulations were carried out in each case for 5,6,7, and 8 years of simulation time, with 100 simulations at each time point. Fits shown are linear for the SSA and logarithmic for leaping. The SSA is shown to result in equivalent or faster execution times in small populations but takes prohibitively long times to simulate at larger population sizes. The particular simulation parameters and leaping threshold used determine the crossover point, typically at 10^3^ − 10^4^ for *x*_thresh_ = 10^2^.

##### Algorithm 2: SSA simulation of duration time SSA_time for set of subpopulations {*X_SSA_*} comprising the full system (SSA_type = pure) or a subsystem (SSA_type = hybrid starting at simulation time *t*)

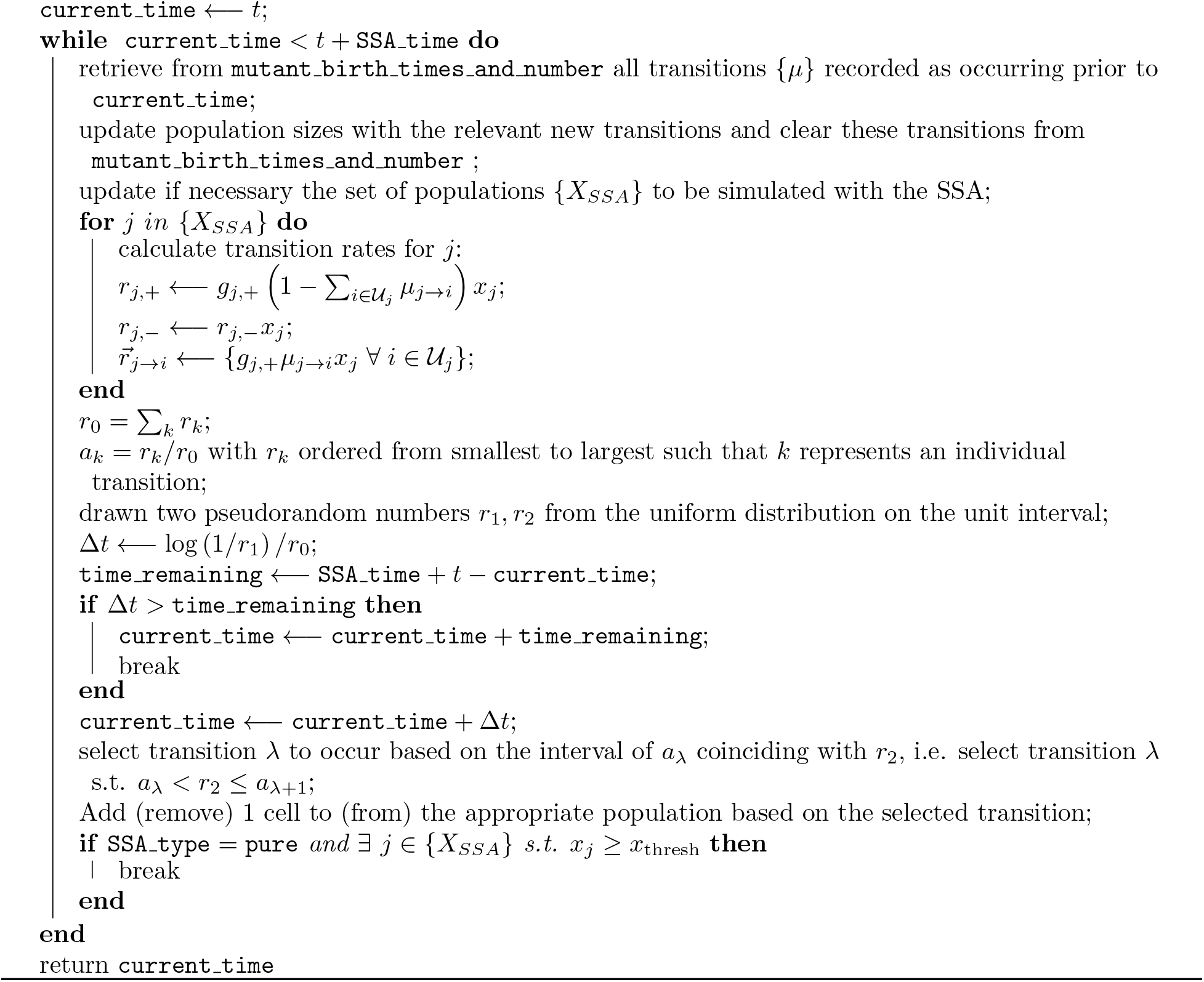

## A Appendix

The sampling variance at any particular step is given by

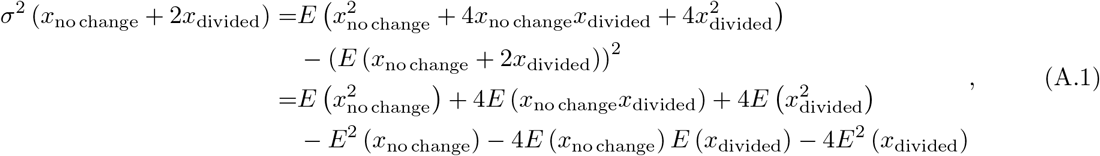

where

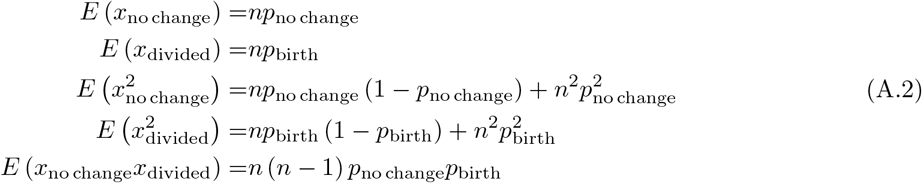

where in the last line we have used that *E* (*X_i_X_j_*) = cov (*X_i_, X_j_*) + *E* (*X_i_*) *E* (*X_j_*). Setting *p*_no change_ = 1 − (*p*_birth_ + *p*_death_), we have that

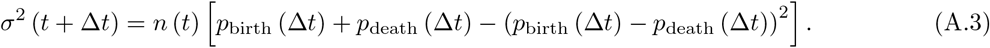

## Footnotes

1 The discussion here does not in principle preclude rate functions with a dependence on *x_j_*, e.g. as in the case of logistic growth; here, however, we restrict ourselves to the linear case.

## Notes

### Competing Interest Statement

The authors have declared no competing interest.

